# Fall Risk Prediction in Multiple Sclerosis Using Postural Sway Measures: A Machine Learning Approach

**DOI:** 10.1101/410704

**Authors:** Ruopeng Sun, Katherine L. Hsieh, Jacob J. Sosnoff

## Abstract

**Background:** Balance impairment affects over 75% of individuals with multiple sclerosis (MS), and leads to an increased risk of falling. Numerous postural sway metrics have been shown to be sensitive to balance impairment and fall risk in individuals with MS. Yet, there are no guidelines concerning the most appropriate postural sway metrics to monitor impairment. This investigation implemented a machine learning approach to assess the accuracy and feature importance of various postural sway metrics to differentiate individuals with MS from healthy controls as a function of physiological fall risk.

**Methods:** This secondary data analysis included 153 participants (50 controls and 103 individuals with MS) who underwent posturography based balance assessment (30s eyes open standing on a force platform) and physiological fall risk assessment (Physiological Profile Assessment - PPA). Participants were further classified into four subgroups based on fall risk: controls (n=50, 64.9 ± 4.9 years old, PPA < 1); low-risk MS (n=34, 54.0 ± 13.1 years old, PPA < 1); moderate-risk MS (n=27, 58.3 ± 8.3 years old, 1 ≤ PPA < 2); high-risk MS (n=42, 56.8 ± 9.7 years old, PPA ≥ 2). Twenty common sway metrics were derived following standard procedures, and subsequently used to train a machine learning algorithm (random forest – RF, with 10-fold cross validation) to predict individuals’ fall risk grouping. The feature importance from the RF algorithms was used to select the strongest sway metric for fall risk prediction.

**Results and Discussion:** The sway-metric based RF classifier had high classification accuracy in discriminating controls from MS individuals (> 86%). Sway sample entropy, a sway regularity metric, was identified as the strongest feature for classification of low-risk MS individuals from healthy controls. Whereas for all other comparisons, mediolateral sway amplitude was identified as the strongest predictor for fall risk groupings. These findings may set the foundation for the development of guidelines for reporting balance impairment in individuals with MS.

## 1. Introduction

Multiple Sclerosis (MS) is a chronic, inflammatory-mediated neurological disorder affecting more than 2.3 million people worldwide[1]. MS is characterized by inflammatory demyelination and axonal damage in the central nervous system[2]. The neuronal damage alters a wide range of cognitive, sensory and motor functions, which contribute to impairment in balance [2]. Balance impairment affects over 75% of persons with MS (PwMS) during the disease course[3], and it is associated with an elevated risk of falls and declines in quality of life[4]. Given its importance, identifying and treating balance impairment is often a focus of MS rehabilitation and research.

Traditionally, functional assessment and self-report questionnaires have been utilized to evaluate balance impairment in PwMS. Functional tests such as Berg Balance Scale (BBS) [5] measures static and dynamic balance ability, whereas self-administered questionnaires such as Activity-Specific Balance Confidence Scale (ABC) [6] and Falls Efficacy Scale - International (FES-I) [7] measure individuals’ balance confidence and fear of falling during daily life. However, these measures rely on subjective scoring and have poor sensitivity due to ceiling or floor effects[8]. On the other hand, instrumented measures through posturography, which utilizes a force platform to quantify Center of Pressure (COP) movement (an indicator for the neuromuscular control of maintaining an upright stance[9]), provides objective and quantitative measures of postural stability and often is considered the gold standard for balance assessment[10]. An advantage of posturography is that it provides precise measurement of movement over fast time scales which affords measurement of subtle alterations in the control of balance.

A number of investigations have analyzed postural impairment in PwMS utilizing posturography. Different variables of postural sway are often utilized because they are thought to reflect different underlying physiological control mechanisms[9, 11]. For example, sway velocity has been reported to be reflective on proprioception function[12], whereas sample entropy, a nonlinear sway regularity/complexity measure, indirectly reflects the ability to adapt to environment [13], such as postural perturbations [14]. However, due to the large number of outcome variables, it is difficult to compare results across investigations. Indeed, a recent systematic review[15] focusing on postural impairment in PwMS identified over 100 different variables as outcome measures of posturography. Furthermore, the choices of measures selected is rarely rationalized and results in a lack of consensus in determine the appropriate measure. Consequently, when measuring balance impairment in PwMS, it is difficult to determine which measure to use in order to provide clinical meaningful insights for treatment planning [15].

In a previous attempt to quantify the diagnostic accuracy of static posturography in fall incidence prediction among PwMS, Prosperini et al. [10] utilized various time domain sway measures (i.e. sway velocity, sway path, and sway area) to identify PwMS at risk of falls. They achieved over 70% of classification accuracy using a stepwise logistic regression and identified COP sway path as the only significant predictor of the fall occurrence. However, they provided minimal rational for choice of COP sway metrics and suggested the need for consensus on which sway measures to use when evaluating balance impairment in PwMS.

One solution to identify appropriate measures to differentiate between PwMS from healthy individuals is through the use of machine learning techniques. Indeed, various machine learning approach have been recently utilized to classify individuals with clinical pathology from controls, such as support vector machine (SVM), random forest (RF), k-nearest neighbor (KNN) and neural network[16-19]. Among those approaches, the RF algorithm[20] has its unique advantage over others as it can not only construct a prediction rule to classify outcomes such as balance impairment, but it can also assess and rank variables with respect to their ability to predict the classification outcome [21]. The RF variable importance is determined by the Gini Importance, which is defined as the times a feature is used to split a node, normalized by the number of samples it splits [21]. The RF classification algorithm has also shown excellent accuracy in discriminating Parkinson’s disease participants based on gait and postural measures [17] and has been used to identify important neuroimaging feature for Alzheimer’s disease diagnosis [22]. For more details on random forests, see [20].

Therefore, the aims of this study were: 1) to identify which postural sway measures differentiate between PwMS and healthy controls, as a function of physiological fall risk; 2) to determine the discriminative ability of postural sway measures for fall risk classification (low, moderate, or high) in PwMS. To achieve these aims, we utilized the random forest algorithm to classify individuals’ fall risk based on a comprehensive set of postural sway parameters that contain time-domain, frequency-domain, and non-linear dynamics measures, and calculated the diagnostic accuracy (sensitivity, specificity, and accuracy) and feature importance of the postural sway measures.

## 2. Methods

This study is secondary analysis of previously published and unpublished data focusing on mobility in MS [23-28]. All data were sampled from baseline assessments prior to any interventions.

### 2.1. Participants

Data from 153 participants (50 healthy controls and 103 individuals with MS) were included in the analysis. The inclusion criteria for MS participants included a previously neurologist-confirmed diagnosis of MS and the ability to stand upright for 30s without aid. Self-reported disability was accessed with the self-reported expanded disability status scale (EDSS_SR_) [29], with higher scores indicating higher functional impairment. Inclusion criteria for healthy controls required no history of neurological or orthopedic pathology that might influence balance or mobility, and the ability to stand upright for 30s without aid. All procedures were approved by the University of Illinois at Urbana-Champaign Institutional Review Board, and all participants provided written informed consent prior to participation.

### 2.2. Procedure

On a single visit, participants were instructed to stand upright for 30s on a force platform (FP4060-05-PT-1000, Bertec Corp, Columbus, OH) with their feet shoulder-width apart and eyes open, fixating at a target 2m away. Individual’s risk of falls was assessed using the short form of the Physiological Profile Assessment[30] (PPA, Neuroscience Research Australia, Sydney), which consist of five validated measures of physiological function (visual contrast sensitivity, proprioception, quadriceps strength, reaction time, and postural sway). The individual fall risk index score was derived from the PPA normative database (Z-scores adjusted) for comparison with population norms, and MS participants were further categorized as low risk (<1), moderate risk (1-2), and high risk (>2) [30, 31].

Participants also provided demographic information and completed questionnaires on balance confidence (ABC or FES-I). The ABC and FES are both validated measures for fear of falling, and have been shown to be highly correlated (r=0.88) [32]. The FES-I contains 16 items scored on a four-point scale (1=not at all concerned to 4= very concerned), which assesses the degree of perceived self-efficacy at avoiding a fall during basic activities of daily living (ADL) [7]. The total score of FES-I ranged from 16-64, with lower score indicating higher perceived self-efficacy at avoiding a fall. The ABC balance scale (a 16-item scale) measures individual’s confidence in maintaining balance while performing ADL [6]. The total score of ABC ranges from 0-100%, and higher score indicate higher confidence in maintaining balance. Due to differences in research procedure across multiple studies, 37 participants only completed the ABC scale, 27 participants only completed the FES-I scale, and 89 participants completed both ABC and FES-I scale. In order to compare the self-reported balance confidence across multiple studies, the FES-I score was converted to a percentage score (0-100%, with higher score indicate higher confidence) similar to the ABC score using the following formula: *FES*_*p*_ = 100% * *abs*(*FES* - 64) /(64 - 16)

The self-reported Balance Confidence was derived as the average of ABC and FES_p_. If only ABC or FES was available, then that available measure was used.

All participants also completed the Berg Balance Scale assessment[5], which consists of 14 physical tasks, such as transfer from sitting to standing position, standing with eyes closed, and picking up an object from the floor, all of which are part of normal daily activities. Each task performance is assigned 0–4 points by a trained personnel to give a total score of 0–56.

### 2.3. Data Processing and feature extraction

COP data were sampled at 1000 Hz and low pass filtered (4th order Butterworth) at 10 Hz for further analysis. A comprehensive set of common postural sway measures that includes time and frequency domain measures and non-linear dynamics were derived from the COP sway data using established procedures [9, 33] and a customized MATLAB program (Mathworks, Inc., Natick, MA): Sway path length (Anterior-Posterior-AP, Medial-Lateral-ML, and Resultant Distance-RD); Mean sway velocity (AP, ML, and RD), 95% confidence ellipse sway area, Sway range (AP, ML direction), Root mean squared sway amplitude (AP, ML); total power, centroidal frequency, frequency dispersion, 95% power frequency; Sample Entropy (AP, ML), and Approximate Entropy (AP, ML). For both Sample Entropy and Approximate Entropy calculation, the input parameter were chosen as m=3, r=0.2, according to Rhea et al. [33].

### 2.4. Model training and performance evaluation

To provide clinical meaningful output for MS fall risk diagnosis, Random Forest algorithms were employed to classify individuals’ fall risk categories (i.e. low, moderate, high) based on postural sway metrics. The RF algorithm is an ensemble of random decision trees (in our case, 1000 trees, chosen by examining the convergence of classification accuracy as additional trees were added [34]), in which the final predicted class for a test example is obtained by combining the predictions of all individual trees. This classifier has been shown to improve the generalization performance of individual decision trees [20].

Binary classification between healthy controls and MS groups (HC vs MS_Low_, HC vs MS_Mod_, HC vs MS_High_, MS_Low_ vs MS_Mod_, MS_Low_ vs MS_High_, MS_Mod_ vs MS_High_) were performed based on sway measures and on clinical balance measures (BBS and Balance Confidence) separately. All classification algorithms were implemented using the open-source machine learning library Scikit-learn in Python [35].

All algorithms were trained and tested through a 10-fold cross validation (CV) scheme[36], and the classification performance was evaluated using standard classification metrics (accuracy, sensitivity, and specificity) derived from the confusion matrix. Briefly, sensitivity was calculated as true-positive/(true-positive + false-negative), specificity as true-negative/(true-negative + false-positive), and accuracy as true-positive + true negative/(true-positive + false-negative + true-negative + false-positive). Since no hyper-parameter tuning was performed in this work (as a proof-of-concept, all algorithms are used in their default settings) and with a limited sample size, data split into training and testing sets was not performed.

In order to identify the importance of each variable in detecting altered balance control in PwMS, a feature selection algorithm was subsequently performed based on the RF classification model. The RF classifier identifies the importance of each input features by calculating the Mean Decrease Impurity (MDI), defined as the number of times a feature is used to split a node, weighted by the number of sample it splits[37]. In other word, if a feature is used multiple times to split a large amount of data, it is identified as a substantial feature.

## 3. Results

Table 1 summarizes the characteristics of the 153 participants (108 Female, 45 Male) in this study. All 50 healthy controls had low fall risk (PPA <1). Participants with MS were further categorized to low risk (n=34, PPA<1), moderate risk (n=27, PPA 1-2), and high risk (n=42, PPA>2) of falls based on PPA. Overall, significant group difference (p<0.05) in age, EDSS, BBS, PPA, and self-reported balance confidence scale were observed within the MS group. MS individuals with increased fall risk also exhibited higher disability (EDSS_SR_), lower balance confidence, and lower functional performance (BBS).

**Table 1.** Participant characteristics. All values were presented with mean and standard deviation, unless otherwise listed. * Significant group difference was observed (p<.05).

|  | Controls | MS<sub>Low</sub> | MS<sub>Mod</sub> | MS<sub>High</sub> |
| --- | --- | --- | --- | --- |
| N (M/F) | 50 (15/35) | 34 (10/24) | 27(7/20) | 42 (13/29) |
| Age* | 64.94 (4.91) | 54.00 (13.12) | 58.26 (8.28) | 56.76 (9.71) |
| EDSS* (Median & IQR) | NA | 4.0 (3.0) | 6.0 (2.0) | 6.0 (0.5) |
| BBS* | 55.12 (2.03) | 50.68 (4.80) | 45.22 (7.82) | 40.05 (8.36) |
| PPA* | -0.08 (0.62) | 0.28 (0.51) | 1.50 (0.31) | 3.01 (0.91) |
| Balance Confidence* | 93.12 (7.05) | 74.67 (19.87) | 58.90 (25.06) | 55.92 (17.19) |

Table 2 provides a summary of the RF classification performance based on sway measures. Sway-based classifier can consistently differentiate healthy controls and MS individuals with classification accuracy of 86.3%, 92.3%, and 89.3% for low, moderate, and high fall risk PwMS, respectively. However, sway-based classifier achieved relatively low accuracy in differentiating MS subgroups (49.5% - 71.2%).

**Table 2.** Diagnostic performance of RF algorithm using posturography measures. ACC-accuracy, SEN-sensitivity, SPC – Specificity

|  | MS<sub>Low</sub> |  |  | MS<sub>Mod</sub> |  |  | MS<sub>High</sub> |  |  |
| --- | --- | --- | --- | --- | --- | --- | --- | --- | --- |
|  | ACC | SEN | SPC | ACC | SEN | SPC | ACC | SEN | SPC |
| Controls | 86.3 | 76.5 | 92.0 | 92.3 | 85.2 | 96.0 | 89.3 | 85.7 | 92.0 |
| MS<sub>Low</sub> |  |  |  | 50.0 | 48.1 | 52.9 | 71.2 | 71.4 | 73.5 |
| MS<sub>Mod</sub> |  |  |  |  |  |  | 49.5 | 66.7 | 22.2 |

Table 3 provides a summary of the RF classification performance based on the BBS and self-reported balance confidence scale. This clinical measure-based classifier achieved lower accuracy in differentiating healthy controls and low risk PwMS (73.5%), but improved the accuracy in differentiating MS subgroups (61.4% - 76.7%), especially in differentiating PwMS with high fall risk among other subgroups (75.9-76.7% in comparison to 49.5-71.2% using only sway measures).

**Table 3.** Diagnostic performance of RF algorithm using BBS and balance confidence measures. ACC-accuracy, SEN-sensitivity, SPC – Specificity

|  | MS<sub>Low</sub> |  |  | MS<sub>Mod</sub> |  |  | MS<sub>High</sub> |  |  |
| --- | --- | --- | --- | --- | --- | --- | --- | --- | --- |
|  | ACC | SEN | SPC | ACC | SEN | SPC | ACC | SEN | SPC |
| Controls | 73.5 | 66.7 | 78.0 | 92.1 | 88.9 | 94.0 | 95.6 | 95.0 | 96.0 |
| MS<sub>Low</sub> |  |  |  | 61.4 | 51.9 | 69.7 | 75.9 | 80.0 | 72.7 |
| MS<sub>Mod</sub> |  |  |  |  |  |  | 76.7 | 87.5 | 59.3 |

The importance of sway measures for differentiating the fall risk categories in PwMS and controls were extracted using the weighted percentage (MDI) of each RF classifier. Figure 1A shows the top-five sway measures for differentiating Low-risk MS individuals from Controls, with Sample Entropy (AP), Sway Range (ML), and Sway Area identified as the top predictors from this model with similar weight (~15%). Figure 1B shows the top sway measures for differentiating Mod-risk MS individuals from Controls. Sway Range (ML) was identified as the governing predictor from this model (62.3% of total importance measure). Figure 1C shows the top sway measures for differentiating High-risk MS individuals from Controls, in which Sway Range (ML), Sway Path Length (ML), and Mean Velocity (ML) were identified as the top predictors from the model (ML sway amplitude account for 71.1% of total importance).

**Figure. 1.**
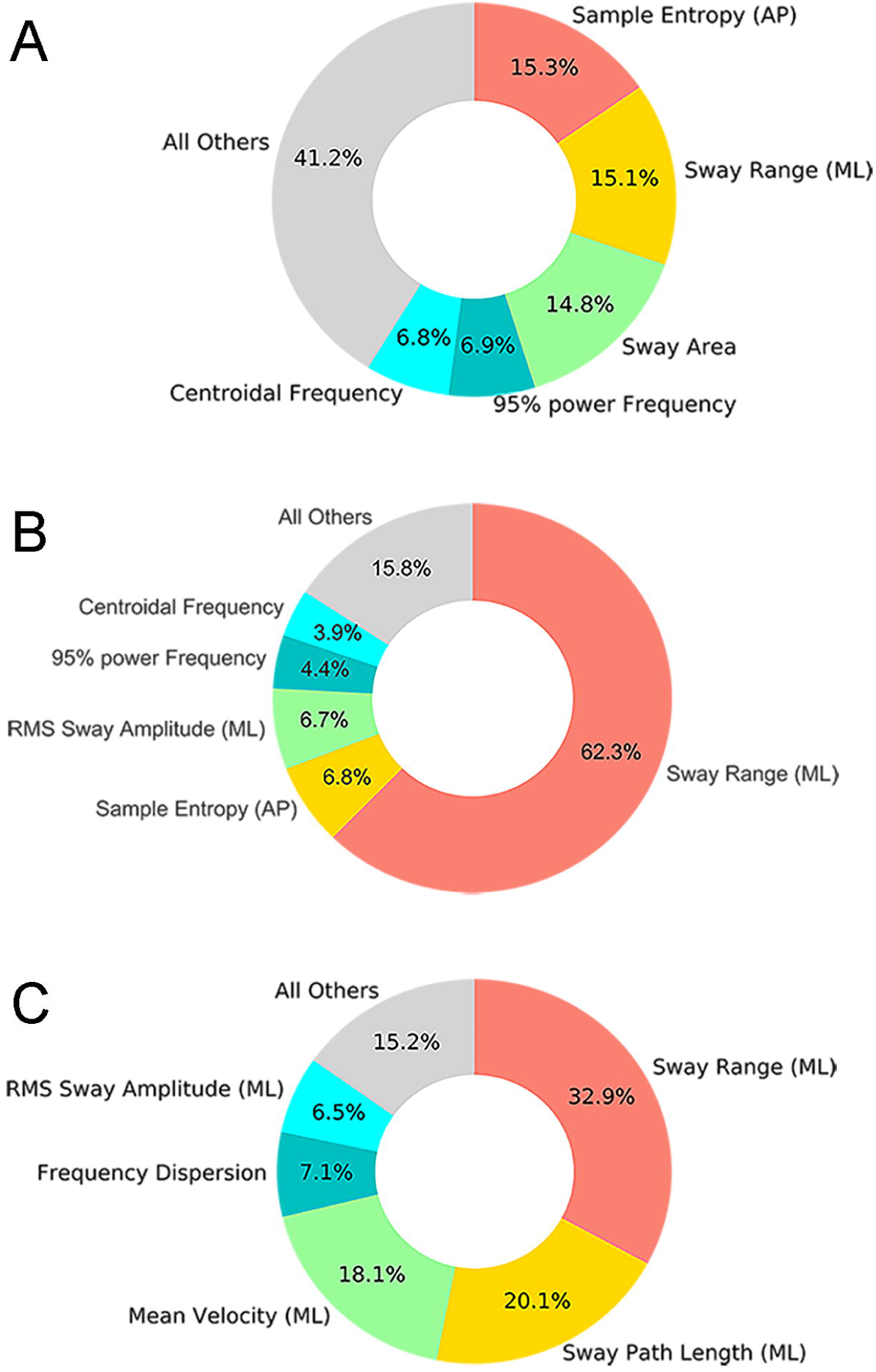
Sway metrics importance as measure by the Mean Decrease Impurity (MDI) of each RF classifier. A) Controls vs MS_Low_; B) Controls vs MS_Mod_; C) Controls vs MS_High_.

## 4. Discussion

This work aimed to identify postural sway parameters that best differentiate between PwMS and healthy controls, as a function of physiological fall risk utilizing a machine learning approach. This work also investigated the discriminative ability of using postural sway measures for fall risk classification in PwMS. Utilizing a machine learning approach (random forest classifier), we demonstrated that postural sway measures can discriminate low-risk PwMS from healthy controls, with over 86% of classification accuracy. In contrast, clinical balance measures (BBS) and self-report balance confidence measures achieved equal to superior discriminative ability in separating MS individuals with moderate and high risk of falls. These findings suggest that posturography measures are sensitive to subtle change in the balance control among MS individuals with minimal fall risk, whereas the balance impairment in moderate and high-risk MS individuals can be assessed with traditional measures with high accuracy.

Overall, the observations further confirm that individuals with MS across the disability spectrum have postural control deficits[15]. The novel contribution of this investigation is identifying distinct COP parameters that are sensitive to balance impairment in MS as a function of fall risk. Indeed, ML sway amplitude parameters including sway range, sway path length, mean sway velocity were the strongest predictors for discriminating moderate and high-risk PwMS from healthy controls. This observation is consistent with Morrison and colleagues[38] who demonstrated that a sample of 22 persons with MS had greater fall risk which coincided with deficits in mediolateral postural control. On the other hand, Sample Entropy, a measure of movement complexity (lower value indicating reduced signal complexity) and potential reduced adaptability to small perturbations [13, 14], was further identified as a key predictor for discriminating low-risk PwMS from healthy controls. This finding is consistent with previous research by Roeing et al. [26] and Huisinga et al. [39] that proposed reduced complexity in postural control as a biomarker for balance impairment in individuals with MS.

Several previous reports have attempted to predict fall risk in PwMS using postural sway measures. For example, Prosperini et al. [10] used time-domain sway measures and logistic regression that achieved over 70% accuracy in discriminating faller from non-faller in PwMS. Furthermore, Kasser et al. [40] utilized a logistic regression to achieve 81% accuracy in discriminating faller and non-faller with differences in the amount of body sway under different sensory conditions (i.e., eyes open to eyes closed). Hoang et al. [31] also utilized logistic regression and achieved 71.2% accuracy (area under the Receiving Operating Characteristic curve) in discriminating faller from non-faller with sway range during eyes closed condition. It is important to note that these previous reports predicted future falls while the current investigation is based on physiological fall risk. Consequently, direct comparisons between investigations may not be appropriate.

One limitation of the present study is that only a single postural control condition was included. Altered sensory test conditions which are common in balance assessments (i.e., eyes closed, standing on compliance surface) were not included in the data analysis. It has been reported that static posturography under altered sensory condition may yield better discriminative ability between PwMS and healthy controls [41], and thus could further improve the classification performance. Another limitation of this work is the relative small sample size (n=30-50 per group), which may limit generalization. It is also important to note that the current observations relate to postural assessment at one point in time. Further research is needed to determine which posturography metrics are most sensitive to rehabilitation intervention. Lastly, the present work did not measure the fall incidence occurrence, thus how the results relate to prediction of future falls is not clear. However, findings from this work may set the foundation for the development of guidelines for accurate reporting of balance impairment in PwMS.

## 5. Conclusion

The current findings highlight the benefits of posturography for balance impairment and fall risk assessment among PwMS, and provide insights on the standardization of metrics. Specifically, static posturography is sensitive for discriminating PwMS with minimal fall risk from healthy controls, with sample entropy measure identified as a core predictor. Whereas for quantification of balance impairment in high-risk MS individuals, medio-lateral sway amplitude is the strongest predictor. Furthermore, clinical balance measures also achieved high discriminative ability in separating MS individuals with moderate and high risk of falls. Therefore, the assessment technique and sway metrics should be based on the target populations, i.e. utilizing posturography (sway entropy) to identify individuals with subtle balance impairment and using clinical balance measures to assess individuals with high fall risk. These findings from this work may set the foundation for the development of guidelines for accurate reporting of balance impairment in PwMS.

## Acknowledgements

The authors would like to thank Dr. Salma Musaad at the Illinois Biostatistics Core of the Interdisciplinary Health Sciences Institute at Illinois for the assistance and feedback in the preparation of this manuscript. Data from this work was collected in previous studies that were funded in part by the National Multiple Sclerosis Society, Consortium of MS Centers, and MC10, Inc.

